# What drives mortality among HIV patients in a conflict setting? A prospective cohort study in the Central African Republic

**DOI:** 10.1101/437103

**Authors:** Thomas Crellen, Charles Ssonko, Turid Piening, Marcel Mbeko Simaleko, Diemer Henri St. Calvaire, Karen Gieger, M. Ruby Siddiqui

## Abstract

**Background:** Provision of antiretroviral therapy (ART) during conflict settings is rarely attempted and little is known about the expected patterns of mortality. The Central African Republic (CAR) continues to have a low coverage of ART despite an estimated 120,000 people living with HIV and 11,000 AIDS-related deaths in 2013. We present results from a cohort in Zemio, Haut-Mboumou prefecture. This region had the highest prevalence of HIV nationally (14.8% in 2010) and was subject to repeated attacks by armed groups on civilians during the observed period.

**Methods:** Conflict from armed groups can impact cohort mortality rates i) directly if HIV patients are victims of armed conflict, or ii) indirectly if population displacement or fear of movement reduces access to ART. Using monthly counts of civilian deaths, injuries and abductions, we estimated the impact of the conflict on patient mortality. We also determine patient-level risk factors for mortality and how this varies with time spent in the cohort. Model-fitting was performed in a Bayesian framework, using generalised-linear models with terms accounting for temporal autocorrelation.

**Results:** Patients were recruited and observed from October 2011 to May 2017. Overall 1631 patients were enrolled, giving 4107 person-years and 148 deaths. Our first model shows that patient mortality did not increase during periods of heightened conflict. The monthly risk (probability) of mortality was markedly higher at the beginning of the program (0.047 in November 2011 [95% credible interval; CrI 0.0078, 0.21]) and had declined greater than ten-fold by the end of the observed period (0.0016 in June 2017 [95% CrI 0.00042, 0.0036]). Our second model shows the risk of mortality for individual patients was highest in the first five months spent in the cohort. Male sex was associated with a higher mortality (odds ratio; OR 1.7 [95% CrI 1.2, 2.8]) along with the severity of opportunistic infections at baseline.

**Conclusions:** Our results show that chronic conflict did not appear to adversely affect rates of mortality in this cohort, and that mortality was driven predominantly by patient specific risk factors. In areas initiating ART for the first time, particular attention should be focussed on stabilising patients with advanced symptoms.

**Funding:** Médecins Sans Frontières

## Introduction

Populations in sub-Saharan Africa with a high burden of HIV/AIDs are often subject to armed conflict. In the absence of humanitarian assistance, people living with HIV (PLHIV) face challenges in accessing treatment or medical services during conflict. Provision of anti-retroviral therapy (ART) during conflict has historically not been prioritised due to a perception that this is too difficult to achieve [1]. In the face of chronic conflict, there was a concern in the humanitarian community that “treatment cannot wait” [2]. The medical organisation Médecins Sans Frontières (MSF) reported results from 22 ART programs in conflict or post-conflict settings in sub-Saharan Africa, finding that patient outcomes were comparable to those in stable resource-limited settings [3]. These findings were also shown in a systematic review and meta-analysis that found mortality and loss to follow-up (LTFU) of patients on ART in conflict settings after twelve months was 9.0% and 8.1% respectively, which are within the range of mortality and LTFU estimates from stable settings [4].

Reports of HIV patient outcomes generally focus on summary statistics after defined intervals (six or twelve months) and are rarely explored as a time series. It is important to consider temporal variation in patient mortality as this has implications for management of patient care and allocation of resources during ART programs. For instance the risk of mortality among patients in sub-Saharan Africa is particularly high in the first three months of ART provision [5, 6] due to individuals seeking treatment at a later stage of disease progression [7]. In the case of studies on the impact of conflict, longitudinal data is particularly important as instability is rarely temporally uniform and often consists of periods of relative calm punctuated by acute violence [2].

The Central African Republic (CAR) is one of the world’s least developed nations [8] and faces a number of challenges in healthcare provision including disruption of services due to instability and civil conflict along with a lack of infrastructure, particularly outside of the capital Bangui. Despite an estimated 120,000 PLHIV, access to ART has been low nationally (13.8% in 2010) and 11,000 AIDS-related deaths in 2013 [9], consequently the mortality rate of 91 deaths per 1000 PLHIV annually is the highest in the world [10].

A nationwide HIV seroprevalence study in 2010 found the Haut-Mbomou prefecture to have the highest prevalence in CAR; 14.8% compared with a prevalence of 5.9% nationally among individuals 15-49 years of age [11]. The health centre in Zemio, Haut-Mboumou prefecture has been supported by MSF since 2010 and an ART cohort was established in October 2011. This was initially a response to an influx of Congolese refugees and internally displaced persons (IDPs), who had fled their homes due to repeated attacks by the Lord’s Resistance Army (LRA) during 2008-10. The region has since faced repeated violent incursions from armed groups [12]. The Zemio HIV treatment program therefore represented an attempt to provide access to ART for a population with a high disease burden in a conflict setting.

Here we explore temporal patterns of mortality among patients in the Zemio HIV cohort over five years in the context of ongoing instability. Conflict from armed groups can impact mortality rates directly if HIV patients are victims of armed conflict, or indirectly if population displacement or fear of movement reduces adherence to ART [13]. We apply generalized-linear models to assess, i) the impact of the conflict measured though civilian death, injuries and abduction, on the risk of mortality in the cohort and, ii) risk factors for mortality at the level of individual patients.

## Methods

We used data from the treatment program’s commencement (18^th^ October 2011) until 31^st^ May 2017. Patients were recruited prospectively throughout the study period and were drawn from the surrounding area, which includes the Bas-Uele province of neighbouring Democratic Republic of Congo along with Congolese refugee camps and IDP camps around Zemio (Figure 1). The target population consists mainly of the local Azande people who work predominantly as subsistence farmers, and Fulani pastoralists who also reside in the region. Patients qualified for HIV testing i) if they volunteered for HIV testing and counselling, ii) were referred to the testing and counselling program after an appointment in Zemio or nearby health facilities, or iii) during the first antenatal visit for pregnant women. Infants testing positive for HIV after birth were also enrolled into the study.

**Figure 1.**
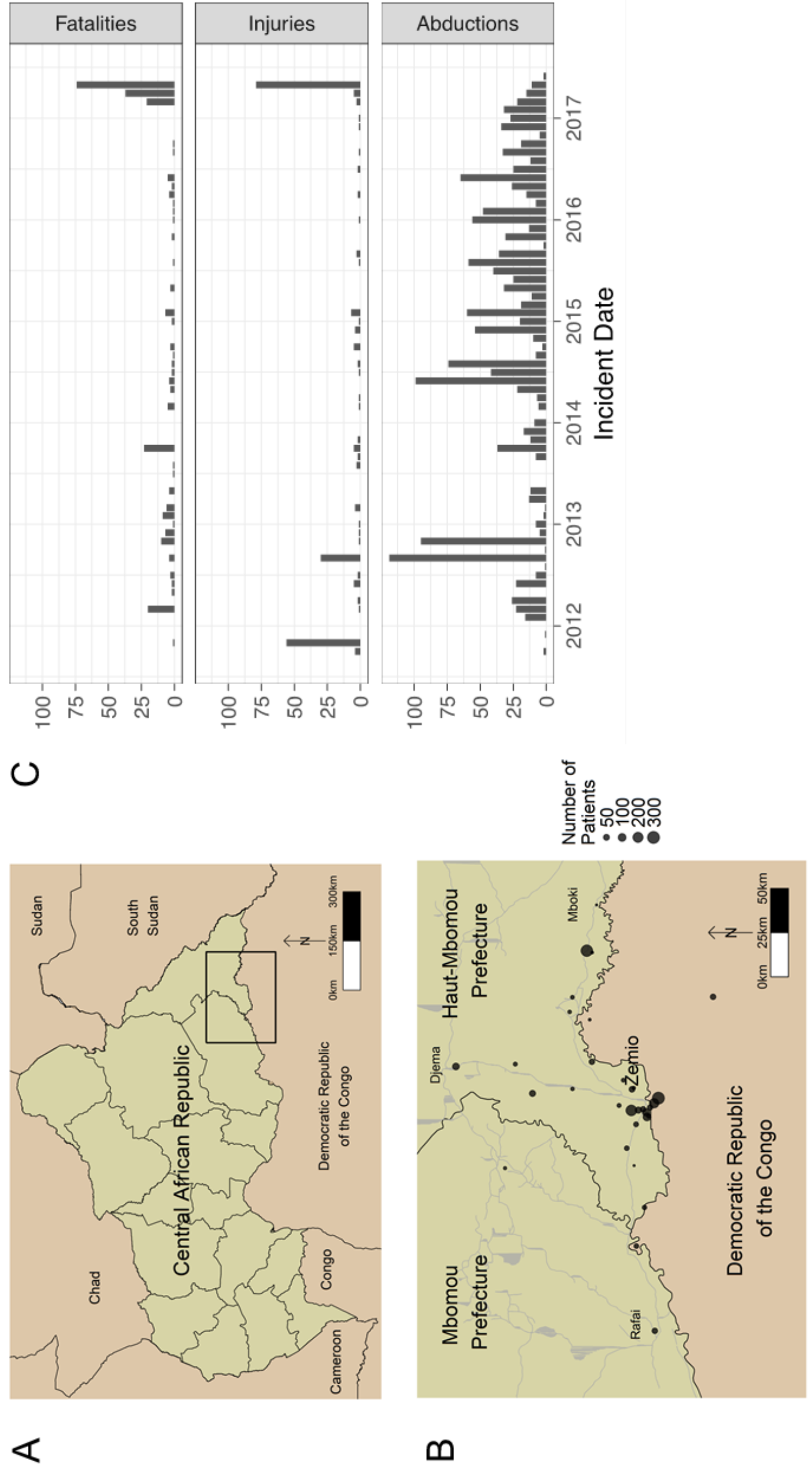
The Central African Republic (CAR) is shown in a regional context (A), with the country divided by prefectures. The study area is indicated by the box in the south east of CAR. The town of Zemio is shown in context (B), within Haut-Mbomou prefecture and adjacent to Mbomou prefecture. Nearby towns where patients were drawn from for the HIV treatment program (Rafai, Djema and Mboki) and the Bas-Uele province of the Democratic Republic of the Congo are also shown. The majority of patients were drawn from around the town of Zemio, including internally displaced person and refugee camps resulting from conflict in 2010 by the Lord’s Resistance Army. Grey lines indicate the road network in the CAR. The number of patients from each location is indicated by the size of the points (*n*=1631). A time series of civilian deaths, injuries and abductions by armed groups in the patient catchment area details the evolution of the conflict in this region from October 2011 – May 2017 (C).

Patients were tested for HIV with i) the Determine™ HIV-1/2 (Alere) antigen/antibody test, ii) Uni-Gold Recombigen (Trinity Biotech) antibody test using whole blood, and iii) ImmunoComb dot ELISA (Biogal). The tests were repeated after three months in the case of discordant results.

The program offered anti-retrovirals (ARVs) to individuals based on WHO guidelines [14]; from 2011-2012 for patients with CD4 <350 cells/μL, from 2013-2016 with <500 cells/μL and for all patients regardless of CD4 count from 2017 (universal test and treat). Patients with CD4 counts above the threshold were given prophylactic treatment against opportunistic infections (OIs) and retested every six months until they were eligible to initiate. Anti-retroviral therapy was offered according to WHO recommendations at the time of entry [15, 16]; first line therapy was typically tenofovir/lamivudine/ efavirenz (TDF/3TC/EFV) combination therapy, second line therapy was zidovudine/lamivudine/ lopinavir/ritonavir (AZT/3TC/LPVr). Patients aged ≤15 years old were treated with paediatric drug formulations. Co-trimoxazole prophylaxis was given to all patients before the initiation of ART and continued during ART. Patients were given appointments at least every three months when ART was provided and clinical signs of OIs were recorded and treated. All diagnostics, treatment and appointments were provided to patients free of charge by MSF.

The Zemio HIV program was mainly nurse driven, an expatriate nurse was present to provide training and guidance to two national staff nurse consultants and *secouristes* (trained lay workers) who provided HIV testing and counseling and psychosocial care to patients. HIV test results were confirmed by a lab technician supported by *secouristes*. A medical doctor was consulted from time to time to provided support on patients with complications. The Zemio team was also supported by the HIV/TB adviser, health adviser and epidemiologist from MSF headquarters. For the supply of ART, MSF ordered ARVs and OI drugs internationally and delivered them to Zemio by plane until 2015, after which supplies were partly ordered through Global Fund in Bangui. An important aspect of the program was community engagement, MSF used 50 community health workers to build acceptance of the program, report on patient outcomes and encourage patients to remain on ART.

Mortality during the study was reported directly if patients died in hospital, or if the patient died in the community this was reported by community health workers. Data was recorded on patient forms and entered into a modified Microsoft Access database during patient enrolment and consultations.

In the first model we consider mortality as a binomial process, with *d* patient deaths from *n* patients at risk with probability *p* of mortality in month *i*.

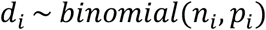

The log-odds of mortality in a given month is a linear function consisting of intercepts (α + γ), explanatory variables (*x*) and coefficients that are estimated by the model (β). The function is constrained between zero and one with the logit link function:

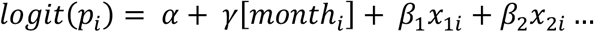

The linear function contains a global intercept (α) and term that varies for each month (γ). The monthly γ is drawn from a multivariate normal distribution, with a vector of zero means and a covariance matrix *K* which specifies the relationship between the probability of mortality on any two months, *i* and *j*, as a function of the length of time between them (*D*_*ij*_). The maximum covariance is specified in *η^2^* and the covariance between months *i* and *j* declines exponentially with the time between them at a rate determined by *ρ*.

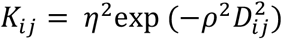

In the second model, death for patient *k* in month *t* (*d_kt_*) is a Bernoulli trial, where *t* is the number of months since entry to the cohort. A term is included that permits variation of the intercept for each month since entry (*γ*) and is drawn from a multivariate normal distribution as described above:

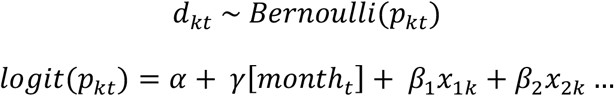

Covariates for model one were the numbers of i) civilian deaths, ii) injuries and iii) abductions per month in the patient catchment area, as recorded by the NGO Invisible Children [12]. Covariates for model two were i) patient’s age at cohort entry, ii) sex and iii) clinical stage of OIs.

Data cleaning and descriptive analysis was performed using R (version 3.4.4) and the models were fitted with Hamiltonian Markov Chain Monte Carlo (MCMC) using Stan (v2.17.3). We assigned vaguely informative normal prior distributions to parameters. Three parallel MCMC chains were run for a minimum of 20,000 iterations including burn-in, and convergence was assessed using the Rubin-Gelman diagnostic and visual inspection.

## Results

A total of 1631 HIV positive patients were recruited into the study, 1147 were female (70.3%) and the median age at first visit was 29 years (interquartile range [IQR] 22, 37). All patients were initiated on co-trimoxazole and the majority were enrolled on ART (*n*=1491, 91.4%). In total there were 148 deaths (9.1%), of which the majority, 121, were among patients on ART. Patients spent a median of 27 months in the cohort (IQR 12, 48) and there were a total of 4107 person-years at risk. We define loss to follow-up as patients that were alive but not present for an appointment >12 months before the end of the program, 183 patients met these criteria. Patients that were CD4 tested on their first visit (*n*=1017) had a median count of 268 CD4 cells/μl (IQR 148, 427), this rose after six months on ART to a median of 403 CD4 cells/μl (IQR 240, 634; *n*=442) and after 12 months to a median of 433 CD4 cells/μl (IQR 264, 636; *n*=321). Patient characteristics are summarised in Table 1.

**Table 1.** Characteristics of the 1631 HIV-positive patients enrolled in the Zemio HIV cohort, Central African Republic from October 2011-May 2017. Interquartile range = IQR, ART= antiretroviral therapy. Loss to follow up is defined as patients not present for an appointment >12 months before the end of the program.

|  |  |
| --- | --- |
| **Total patients enrolled in the Zemio HIV cohort** | 1631 (4107 person-years at risk) |
| **Patients enrolled on ART** | 1491/1631 (91.4%) |
| **Female patients** | 1147/1631 (70.3%) |
| **Median age at first appointment (IQR)** | 29 years (22, 37) |
| **Deaths** | 148 |
| **Loss to follow up** | 183 |
| **CD4 cell count/μl , baseline (IQR), *n*=1017** | 268 (148, 427) |
| **CD4 cell count/μl, 6 months (IQR), *n*=442** | 403 (240, 634) |
| **CD4 cell count/μl, 12 months (IQR), *n*=321** | 433 (264, 636) |

Modelling patient mortality as a series of binomial trials, we estimated the probability of mortality for each month of the study (Table 2). The results show a secular trend, whereby the risk of mortality was high in the initial months of the study and subsequently fell. In November 2011 the median probability of mortality from the marginalised posterior distribution was 0.049 (95% CrI 0.0078, 0.22), this declined throughout 2012 until reaching a steady state by the start of 2013 (Figure 2 panel A). By the end of the observed period in May 2017 the median probability of mortality had fallen greater than 10-fold to 0.0017 (95% CrI 0.00043, 0.0036).

**Table 2.** Counts of patients at risk and deaths over the first 12 complete months of the Zemio HIV cohort, Central African Republic (November 2011-October 2012). The posterior predicted values from model one are shown, which estimate the risk of mortality while accounting for temporal autocorrelation and the impact of covariates (monthly counts of civilian fatalities, injuries and abductions). We present the posterior median, along with 80% and 95% credible intervals. The results show a continuous decline in the risk of mortality over time. The values for all study months are presented in Figure 2 panel A.

| **Month** | **At Risk** | **Deaths** | **Risk of mortality (posterior median)** | **Risk of mortality (95% credible interval)** |
| --- | --- | --- | --- | --- |
| November 2011 | 5 | 1 | 0.049 | 0.0078, 0.215 |
| December 2011 | 22 | 2 | 0.031 | 0.013, 0.079 |
| January 2012 | 37 | 2 | 0.029 | 0.013, 0.063 |
| February 2012 | 58 | 1 | 0.024 | 0.012, 0.047 |
| March 2012 | 97 | 2 | 0.021 | 0.0089, 0.044 |
| April 2012 | 133 | 3 | 0.019 | 0.010, 0.030 |
| May 2012 | 151 | 4 | 0.018 | 0.0098, 0.029 |
| June 2012 | 173 | 1 | 0.014 | 0.0080, 0.022 |
| July 2012 | 189 | 1 | 0.016 | 0.0073, 0.020 |
| August 2012 | 224 | 2 | 0.011 | 0.0065, 0.018 |
| September 2012 | 253 | 4 | 0.0076 | 0.0026, 0.019 |
| October 2012 | 257 | 3 | 0.0087 | 0.0049, 0.014 |

**Figure 2.**
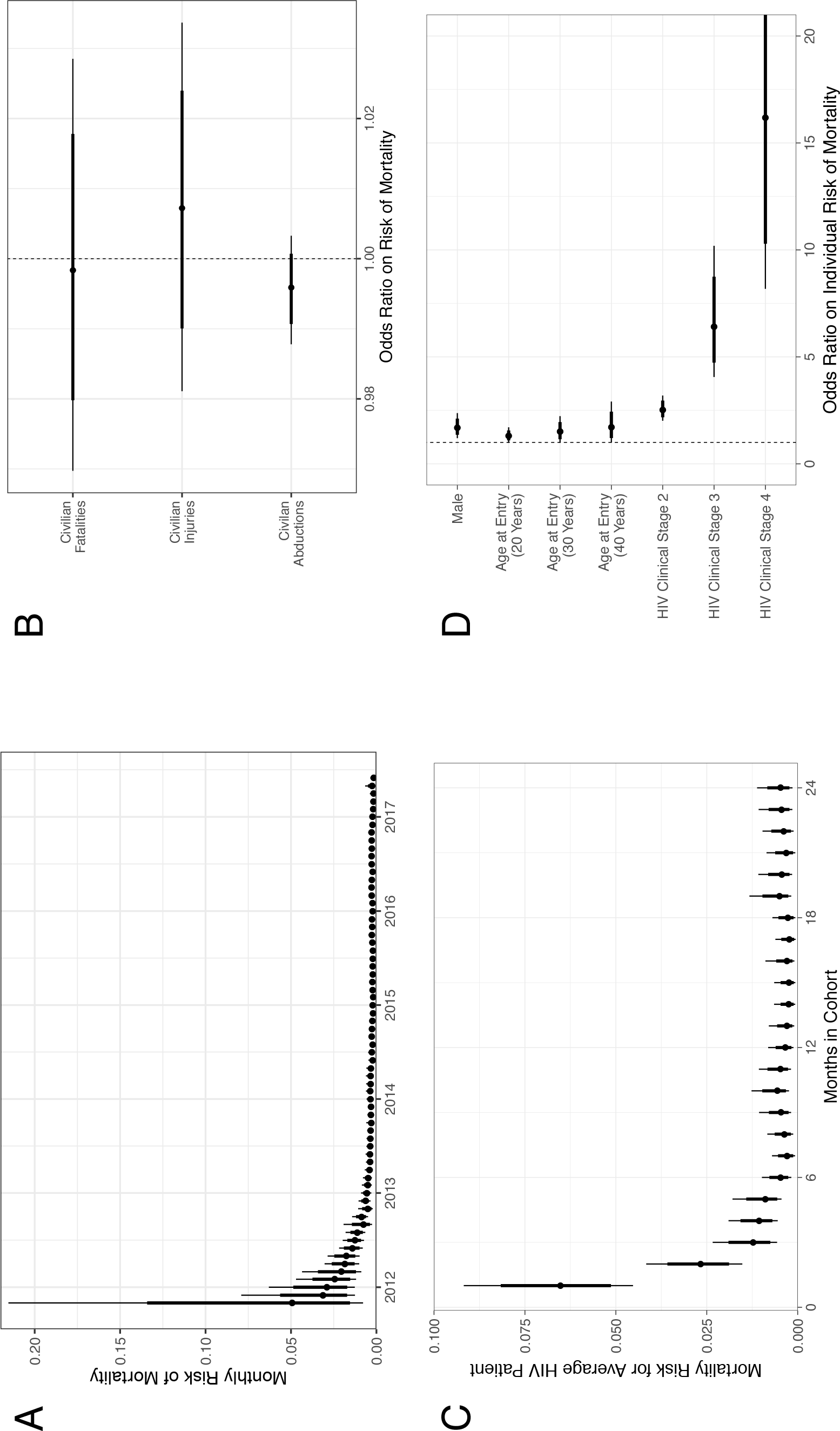
The predictive posterior distributions of the risk of mortality among patients in the Zemio HIV cohort, Central African Republic are shown from models one and two along with the odds ratios of covariates. In all panels, the dot shows the posterior median, and the thick and thin lines show the 80% and 95% credible intervals respectively. The risk of mortality by study month from model one is shown in (A). The corresponding number of deaths and patients at risk for the first 12 months are shown in Table 1. From the odds ratios posterior distributions of model one covariates in panel (B) (unit increase in civilian fatalities, injuries and abductions per month; counts are shown in Figure 1 panel C), the mass of these distributions is centred around one implying that the covariates have no effect on the per-month risk of mortality in the cohort. The estimates of these odds ratios for ten civilian fatalities / injuries / abductions per month are presented in the Results. The estimated risk of mortality by months spent in the cohort are shown in panel (C) (values for the first 12 months are given in Table 2). Odds ratios of covariates from model two are shown in panel D and values are given in the Results. Male sex, higher age at first visit and more severe

From the first model we obtained estimates of the impact of civilian fatalities, injuries and abductions on patient mortality in the cohort; the posterior distributions of odds ratios are centred around one for all three covariates (Figure 2 panel B). As these are continuous variables, we scaled the model coefficients to estimate the impact of 10 community cases of i) fatalities (OR 0.98 [95% CrI 0.74, 1.3]), ii) injuries (OR 1.1 [95% CrI 0.83, 1.4]) or iii) abductions (OR 0.960 [95% CrI 0.88, 1.0]) on cohort mortality. As the mass of the three posterior distributions are centred on one, our model indicates that periods of heightened instability were not associated with higher HIV patient mortality.

In the second model we assessed patient-level covariates for the risk of mortality each month the patient was at risk, with an intercept that varied by months spent in the cohort. Three patients with missing covariates were dropped, leaving 1628 patients with 49,285 person-months at risk. The risk of mortality is shown for a patient with average covariates over the first 24 months (Figure 2 panel C). This risk is highest in the first month (0.065 [95% CrI 0.045, 0.092]) and declines over the first five months to reach a steady state from six months (0.0047 [95% CrI 0.0018, 0.0098]) onwards.

The covariates from the second model estimate that male sex is a risk factor for patient mortality (OR 1.7 [95% CrI 1.2, 2.4]). Older patients at cohort entry show a modest increase in the odds of mortality; i) 20 years at entry (OR 1.3 [95% CrI 1.0, 1.7]), ii) 30 years at entry (OR 1.5 [95% CrI 1.0, 2.2]) and iii) 40 years at entry (OR 1.7 [95% CrI 1.0, 2.9]). The strongest effect, predictably, is HIV clinical staging of OIs at entry. Compared with patients with clinical stage one OIs, the odds of mortality are higher for i) clinical stage two OIs (OR 2.5 [95% CrI 2.0, 3.0), ii) clinical stage three OIs (OR 6.4 [95% CrI 4.1, 10.2], and iii) clinical stage four OIs (OR 16.2 [95% CrI 8.2, 32.5]).

## Discussion

The time series of civilian fatalities, injuries and abductions by numerous armed groups shows the evolution of a “chronic conflict” (Figure 1, panel 3). There was a high underlying rate of civilian abductions throughout the period, predominantly by the LRA. Civilian mortality before 2017 was due mainly to attacks by the LRA or unidentified armed groups on civilians. During 2017 the region became embroiled in a larger national conflict between Christian Anti-Balaka and Muslim Ex-Seleka militias who clashed in the patient catchment area of Mboumou / Haut-Mboumou. Road blocks by the LRA, in addition to a lack of public transport made it challenging for patients to travel to Zemio for appointments or ART collection. Armed robberies of the staff compound were also a frequent occurrence and fighting inside Zemio led to expatriate MSF staff being evacuated twice over the study period[17].

Despite this instability, the pattern of mortality among HIV patients is similar to that observed in longitudinal studies in non-conflict settings, where mortality rates are largely driven by the clinical severity of OIs at baseline [18]. Furthermore, the relatively low numbers lost to follow-up and the recovery of CD4-cell counts suggests that the Zemio HIV program was successful in retaining patients on ART. Mortality was high at the beginning of the program as ART had been absent in this region and a high proportion of patients had advanced symptoms. Of the patients that enrolled in the first three months of the program 36/40 (90.0%) were diagnosed with clinical stage three or four OIs, compared with 14/46 (30.4%) patients that enrolled in the last three months of the program.

The Zemio HIV cohort was transitioned to a community-based model of care in 2017 in order to reduce operating costs, personnel requirements, and ensure the future sustainability of the program. Community models have been shown in low-resource settings to deliver outcomes for patients that are comparable to provider-based care [19, 20]. This transition was interrupted when fighting broke out again in the town of Zemio in July 2017 leading MSF to suspend its operations. The community model of care was continued from December 2017 onwards and MSF continue to monitor the impact of conflict on mortality and patient outcomes.

We have shown that conflict is not a barrier to the provision of ART and that, even in the most challenging environments, HIV patients can adhere to treatment and show favourable outcomes.

## Acknowledgements

The authors would like to thank the members of the Zemio HIV team for their hard work, professionalism and dedication to patients throughout the program: Mr Peeters Muntulongo, Mr Francois d’Assis Koyassama (Nurses), Mr Romuald Gbiakota (Auxiliary Nurse), Mr Joseph Mbolingbagbe, Ms Francisca Zounga-Awo and Mr Martin Gbaboundou (Counsellors), Mr Alain Mbolihinipai, Mr Jacques Koumbodigui (Pharmacists), Mr Hassan Mahamat (Data Encoder) and Mr Igor Mokopa (Laboratory Technician). We would also like to thank Ms Nastasia Stipo and Mr Paul Ronan at Invisible Children for sharing their data on conflict in the Zemio region. In addition, we thank Prof. Ben S. Cooper, Dr. Mo Yin and Dr. Cono Ariti for their valuable comments on the manuscript.

